# GOAT: Deep learning-enhanced Generalized Organoid Annotation Tool

**DOI:** 10.1101/2022.09.06.506648

**Authors:** Jan P. Bremer, Martin E. Baumdick, Marius S. Knorr, Lucy H.M. Wegner, Jasmin Wesche, Ana Jordan-Paiz, Johannes M. Jung, Andrew J. Highton, Julia Jäger, Ole Hinrichs, Sebastien Brias, Jennifer Niersch, Luisa Müller, Renée R.C.E. Schreurs, Tobias Koyro, Sebastian Löbl, Leonore Mensching, Leonie Konczalla, Annika Niehrs, Florian W. R. Vondran, Christoph Schramm, Angelique Hölzemer, Karl Oldhafer, Ingo Königs, Stefan Kluge, Daniel Perez, Konrad Reinshagen, Steven T. Pals, Nicola Gagliani, Sander P. Joosten, Maya Topf, Marcus Altfeld, Madeleine J. Bunders

## Abstract

Organoids have emerged as a powerful technology to investigate human development, model diseases and for drug discovery. However, analysis tools to rapidly and reproducibly quantify organoid parameters from microscopy images are lacking. We developed a deep-learning based generalized organoid annotation tool (GOAT) using instance segmentation with pixel-level identification of organoids to quantify advanced organoid features. Using a multicentric dataset, including multiple organoid systems (e.g. liver, intestine, tumor, lung), we demonstrate generalization of the tool to annotate a diverse range of organoids generated in different laboratories and high performance in comparison to previously published methods. In sum, GOAT provides fast and unbiased quantification of organoid experiments to accelerate organoid research and facilitates novel high-throughput applications.

## Main

Organoids are multicellular 3D structures generated from human tissue cells or induced pluripotent stem cells (iPSCs), mimicking human organs *in vitro*^1–3^. Organoids recapitulate organ-specific cellular functioning^4–6^ and are emerging as powerful systems to model human diseases and perform drug discovery. Furthermore, cells in organoids share gene expression signatures of the tissue of the patient, thereby opening up new strategies for personalized medicine and regenerative therapeutic approaches. The unique opportunities the organoid technology offers for medicine has resulted in an exponential increase of organoid studies for fundamental and clinical research^7,8^. However, analyses of organoid experiments, including growth kinetics and additional organ-specific characteristics, currently rely on subjective visual interpretation of microscopy images or time-consuming annotation tools with manual editing of parameters, introducing variability between measurements performed by different researchers and laboratories^9^. Furthermore, precise monitoring of organoid cultures requires deeper phenotypic characterization of organoids, including assessment of organoid shape, level of budding, density and homogeneity. Although advances have been made in the automated annotation of organoids, these tools are limited by requirements of additional configurations, lack of generalization and robustness, and use of rudimentary annotation methods, such as bounding boxes^10,11^. Taken together, to achieve the potential that organoid systems offer for biomedical research, fast and standardized annotation tools of organoid systems are needed.

Progress in computer vision has advanced the field of object recognition within images, including object detection^12,13^, semantic segmentation^14^ and more recently instance segmentation^15^ with neural networks. However, deep neural networks are prone to overfitting and lack performance on variable datasets. To address these challenges and advance robust organoid studies, we developed a neural-network based tool for generalized organoid annotation (GOAT (Generalized Organoid Annotation Tool)) and evaluated it on a diverse, multicentric dataset consisting of more than 90,000 instance mask annotated six organoid system derived from five different tissues and distributed over 729 brightfield images.

## Results

### Mask R-CNN

GOAT (Generalized Organoid Annotation Tool) is a quantification tool for brightfield microscopy images of organoids (**Fig. 1a**). It was built upon the Mask R-CNN^15^ neural network architecture, utilizing a ResNet-34-FPN^16,17^ backbone and briefly consisting of two sections **(Fig. 1b)**. In the first section, a stack of convolutional layers, including the backbone, produces feature maps that identify regions of interest and thus serves as a region proposal network. In the second section, from these regions of interest, bounding boxes around object instances, as well as classes of these object instances, are determined. Simultaneously, feature vectors from the proposed region are fed into another branch of the same neural network that is configured as a segmentation network. This second branch determines the mask that identifies pixels belonging to each instance of an organoid. (For further details on training see Methods). These instance masks are used to quantify the number and size of organoids. The pixel-level annotation is critical to separate and identify overlapping organoids, which is a common feature of organoid cultures and furthermore allows for the quantification of additional features, such as organoid shape and optic density. This is not achieved by other deep learning approaches, including object detection, which lacks pixel-level precision or semantic segmentation that is not able to separate overlapping organoids (**Fig. 1c**). Taken together, GOAT, using the Mask R-CNN architecture, yields identification and precise phenotyping of organoids in microscopy images.

**Fig. 1:**
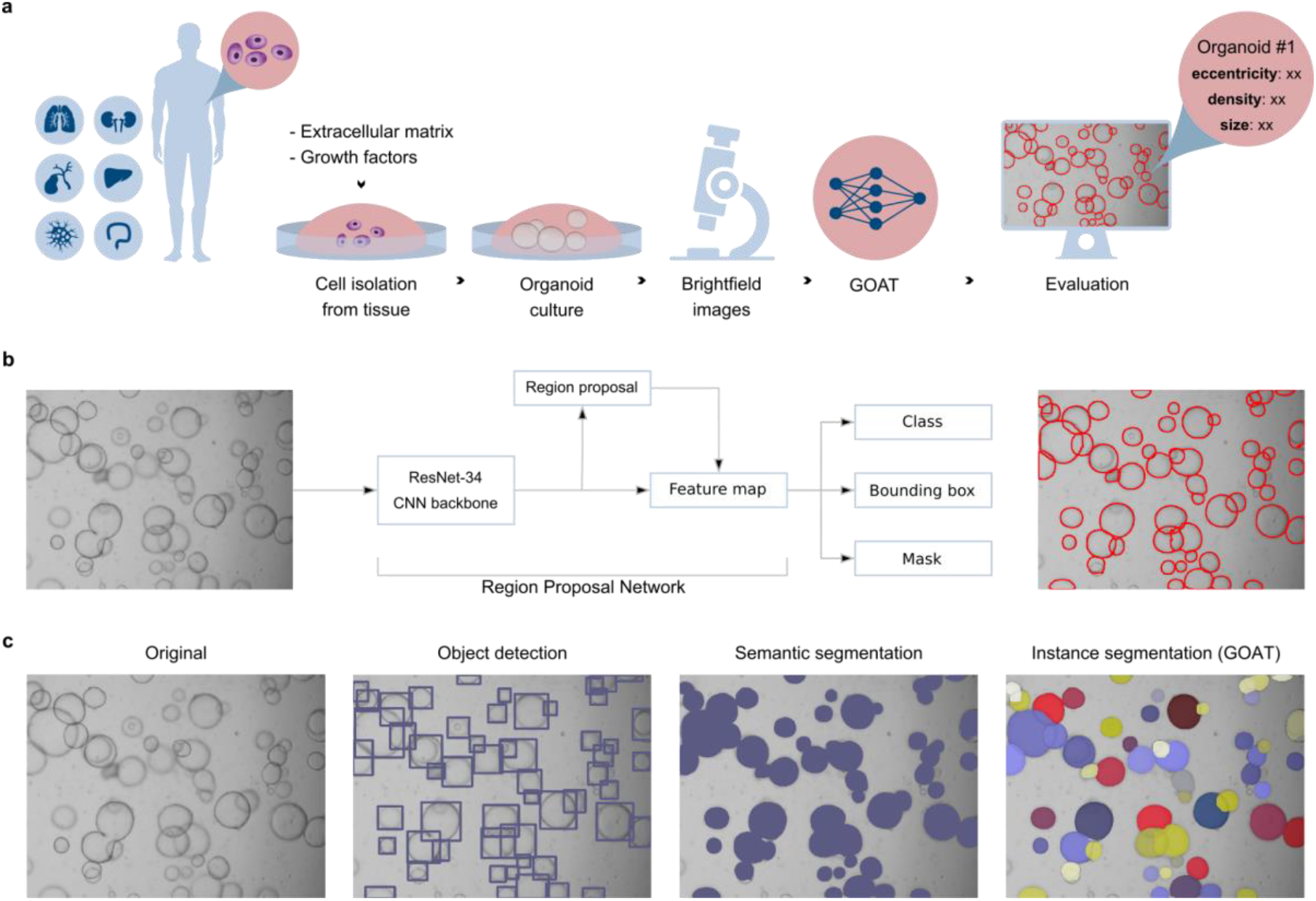
Development of GOAT for annotation of organoid cultures. **a**, Schematic overview to generate organoids from human organs, imaging of organoid cultures and annotation in brightfield images with GOAT. **b**, Schematic representation of the model architecture of GOAT. GOAT was built upon the Mask R-CNN architecture with a Resnet-34 backbone, an extension to the Faster R-CNN architecture. The model in xthe first step performs bounding box prediction, followed by a prediction of a semantic segmentation mask for every bounding box, after which the instance segmentation mask is generated. For every instance segmentation mask a class is predicted, which corresponds to the probability of an instance segmentation to correctly label an organoid and is used to threshold the certainty of the model to make a prediction. **c**, Overview of different deep learning based approaches to annotate organoids in microscopy images, showing the original image, organoids identified with object detection, semantic segmentation and instance segmentation. Instance segmentation allows for pixel precise annotation of overlapping organoids, which can not be achieved with object detection and semantic segmentation.

### Organoid generation and data collection

GOAT was trained on a dataset containing microscopy images of organoid cultures from different tissues, including intestines, colorectal cancer (CRC), bronchoalveolar lavages, liver resections and urine sediment cells, which were generated and cultured as previously described^2,4,5,18–20^. In brief, cells were isolated from tissues or biological fluids and embedded in a 3D scaffold, followed by culture in an organ-adapted medium containing tissue-specific niche factors to promote cell proliferation. Images of intestinal, airway, hepatocyte, and cholangiocyte organoids as well as tubuloids were acquired with brightfield microscopy at the Leibniz Institute of Virology (referred to as Lab (1)). Additional microscopy images obtained at three additional laboratories (Lab (2), Lab (3), Lab (4)) of intestinal organoids and CRC tumoroids were included to increase variability of images of organoid cultures. In sum, a large and diverse dataset of 729 images from six different organoid systems and generated in four laboratories, acquired with five different types of microscopes were used to train and validate GOAT.

### Annotation procedure of organoids

The annotation process was performed by eleven researchers with experience in organoid research. To create instance segmentation ground-truth labels in the first phase, object detection bounding boxes were annotated. These were then cropped and manually segmented in the second phase. Both phases were accelerated using pseudo labeling with manual rechecking. The annotation procedure resulted in a dataset consisting of 729 images containing 90,210 annotated organoids. Subsequently the annotated dataset was split into 488 images for training (67%), 93 images for validation (13%), 12 images for the threshold calculation (2%) and 136 images included in the testing data (19%). The training, validation and threshold calculation datasets consisted of images from Lab (1) of intestinal, airway and hepatic organoid cultures. The testing dataset consisted of organoids from all four laboratories, including intestinal, airway, hepatic and cholangiocyte organoids, tumoroids and tubuloids **(Supplementary Table 1)**.

### Training and validation of GOAT

Following the annotation of organoids within the available images, the Mask R-CNN neural network was trained on the instance segmentation masks using the training data of 488 images. The final neural network weights were selected to be the model at the epoch with the lowest loss on validation data. The training and validation process allowed to derive the final neural network weights at epoch 114 of the training process for GOAT (for details see Methods and GitHub Repository).

### Testing of GOAT

Following training, GOAT was tested on the testing dataset **(Fig. 2)**, generating metrics to evaluate the model’s performance. The main measurement of the performance of GOAT was the mean F1-score at a confidence threshold of 0.9 at an intersection over union (IoU) threshold of 0.50 (F1-50)^21,22^. The F1-50 score is the harmonic-mean of precision and recall and ranges from zero (worst) to one (best). The confidence threshold was chosen by computing the F1 score for incremental threshold values (steps of 0.05) on the threshold calculation dataset. The highest F1-50 score of 0.760 was detected at a confidence threshold of 0.9, which was subsequently used for all further analysis **(Fig. 2b)**. The IoU threshold is commonly selected is 0.50^23^ and results are shown here in **Fig. 2**. In addition, F1 scores (F1-10, …, F1-90) were calculated for different IoU thresholds (**Supplementary Fig. 1a & Supplementary Table 2)**. In sum, the F1-score decreased with higher IoU thresholds, as predictions must have a higher IoU with a ground truth mask to be considered a (true positive) TP **(Fig. 2c)** (For details see Methods and for other performance metrics see **Supplementary Fig. 1a & Supplementary Table 2).**

After the confidence threshold was determined, GOAT was tested on intestinal (mean F1-50: 0.80), airway (0.51) and hepatocyte organoids (0.67) (**Fig. 2d**), showing highest F1-50 on intestinal organoids. To assess generalization of the model to detect organoids generated from other organs, GOAT was tested on images of tubuloids derived from urine and cholangiocyte organoids derived from liver tissue from Lab (1) (**Fig. 2d**). Similar performance of the neural network to detect cholangiocyte organoids and tubuloids with F1-50 scores of 0.77 and 0.78, respectively, was observed. Finally, to determine the reproducibility and robustness of GOAT, the performance on images of intestinal organoids and CRC tumoroids generated in two other laboratories and taken with different microscopes was evaluated (**Fig. 2d**). The performance to detect intestinal organoids from Lab (3) was not significantly different with a F1-50 score of 0.66 (t(31) = 1.41, *P* = 0.17) compared to Lab (1) (F1-50: 0.80), (**Fig. 2d**) and only moderately reduced for organoids from Lab (2) with an F1-50 score of 0.60 (t(46) = 2.97, *P* < 0.01) (**Fig. 2d**). The detection of tumoroids with F1-50 scores of 0.74 (t(17) = 0.72, *P* = 0.48) was similar compared to intestinal organoids from Lab (1) (**Fig. 2d**). In sum, these assessments showed high performance and generalization for organoid systems generated from different organs and laboratories, demonstrating that GOAT can be used to detect and annotate a broad range of organoid systems cultured and analyzed in different laboratories.

**Fig. 2:**
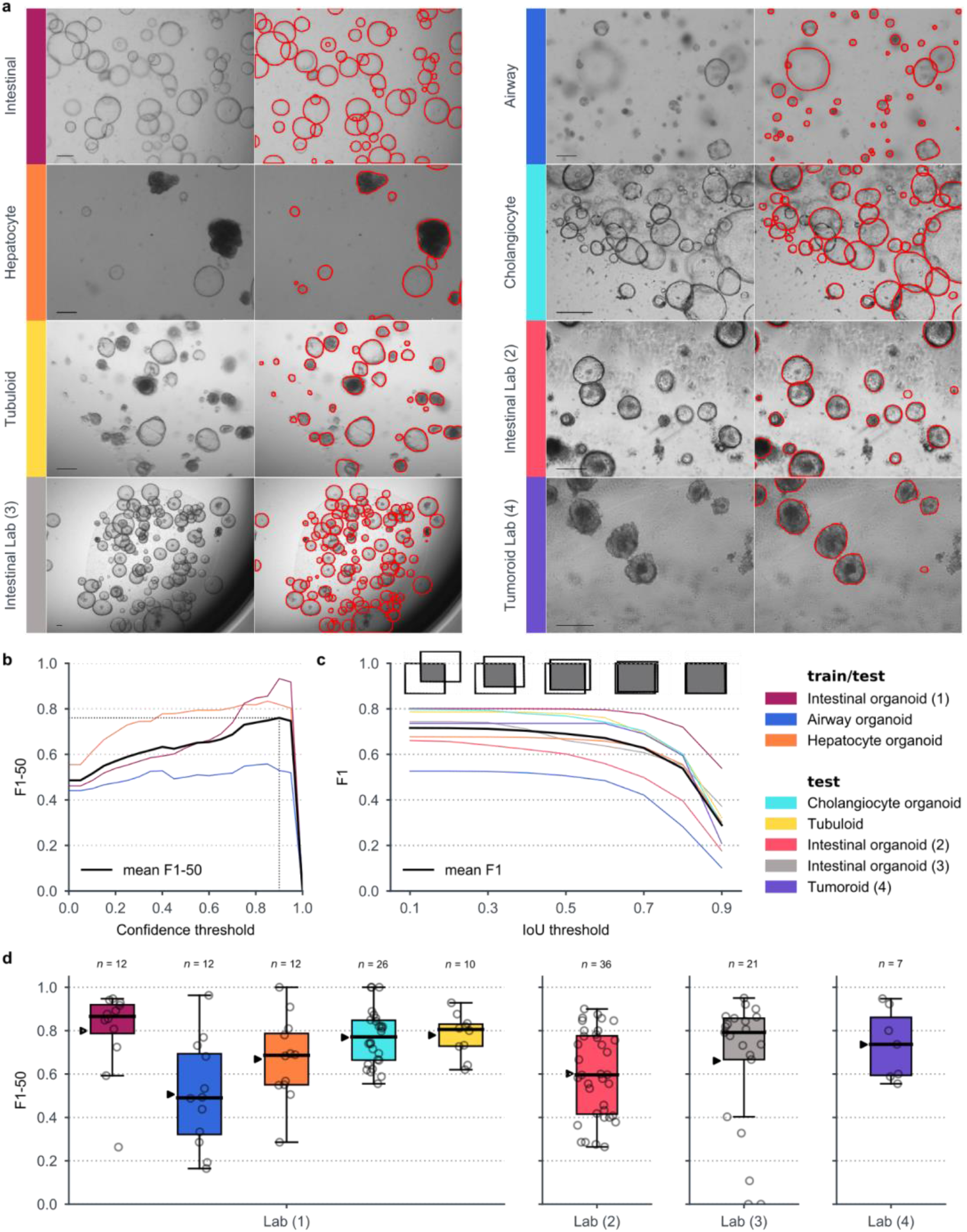
GOAT performance metrics. **a**, Representative images of multiple organoid systems (left) and the corresponding instance segmentation prediction (right), showing intestinal, hepatocyte, airway and cholangiocyte organoids, tubuloids and tumoroids. The instance masks predicted with GOAT are outlined in red. All scale bars represent 250 μm (length variations depends on different microscope settings across images) **b**, F1 scores at different confidence thresholds at an intersection over union (IoU) threshold of 0.5 on the threshold calculation dataset consisting of intestinal (red), airway (blue) and hepatocyte (orange) organoids for determining a confidence threshold for subsequent calculations. The maximal F1-50 score for the mean (black) of the three organoid systems and corresponding confidence score are indicated with the gray dotted line. A maximum F1-50 score of 0.746 was determined at a confidence threshold of 0.9 (for details see Methods). **c**, F1 iterated over different IoU thresholds on the testing dataset at a confidence threshold of 0.9. The mean F1-score is indicated with black. At lower IoU thresholds more predicted masks with smaller overlapping areas with ground truth masks are considered true positives and therefore F1 increases with lower IoU thresholds **d**, F1-50 boxplot for different organoid systems and grouped by the laboratory the experiments were performed. Each dot represents one image. Medians with interquartile ranges are shown. For comparisons, mean F1-50 is marked with a black triangle for each organoid system and laboratory.

### Comparison of GOAT to other organoid annotation tools

Recently, we used organoid systems to study the regulation of human gut development by CD4^+^ T cells^18^. These experiments were performed at Lab (2), the Amsterdam University Medical Center (AUMC). The process of organoid annotation in 319 images (subsequently referred to as gut development (GD) dataset) required the researcher more than two weeks of full-time annotation in the initial study. The annotation was performed with ImageJ, currently the most common method to annotate organoids. In contrast, the analysis of the same dataset by GOAT required 159 seconds (∼2.0 images per second). To assess GOATs applicability against this frequently used method annotations by GOAT and manual annotation with ImageJ were compared (**Fig. 3a, b**). The GD dataset included images from two different microscopes; microscope 1: Leica DM6 wide-field upright microscope; microscope 2: Leica DM6000 wide-field upright microscope. Images taken with microscope 2 showed more organoids out of focus. Organoids out of focus were omitted in the annotation by ImageJ in the initial study. The confidence threshold of predictions for GOAT was kept at 0.9, as described above (**Fig. 2b**). Data obtained by manual annotation and GOAT showed a low mean absolute error and high correlation between the number of organoids predicted by GOAT and the number counted by the researcher, with minimal variability between the two imaging tools (**Fig. 3a**; microscope 1: pearson coefficient = 0.699; *P* < 0.001, *n* = 140 images; microscope 2: pearson coefficient=0.892, *P* < 0.001; *n* = 179 images). Taken together, GOAT produced fast results and low variability between imaging devices compared to manual annotation.

**Fig. 3:**
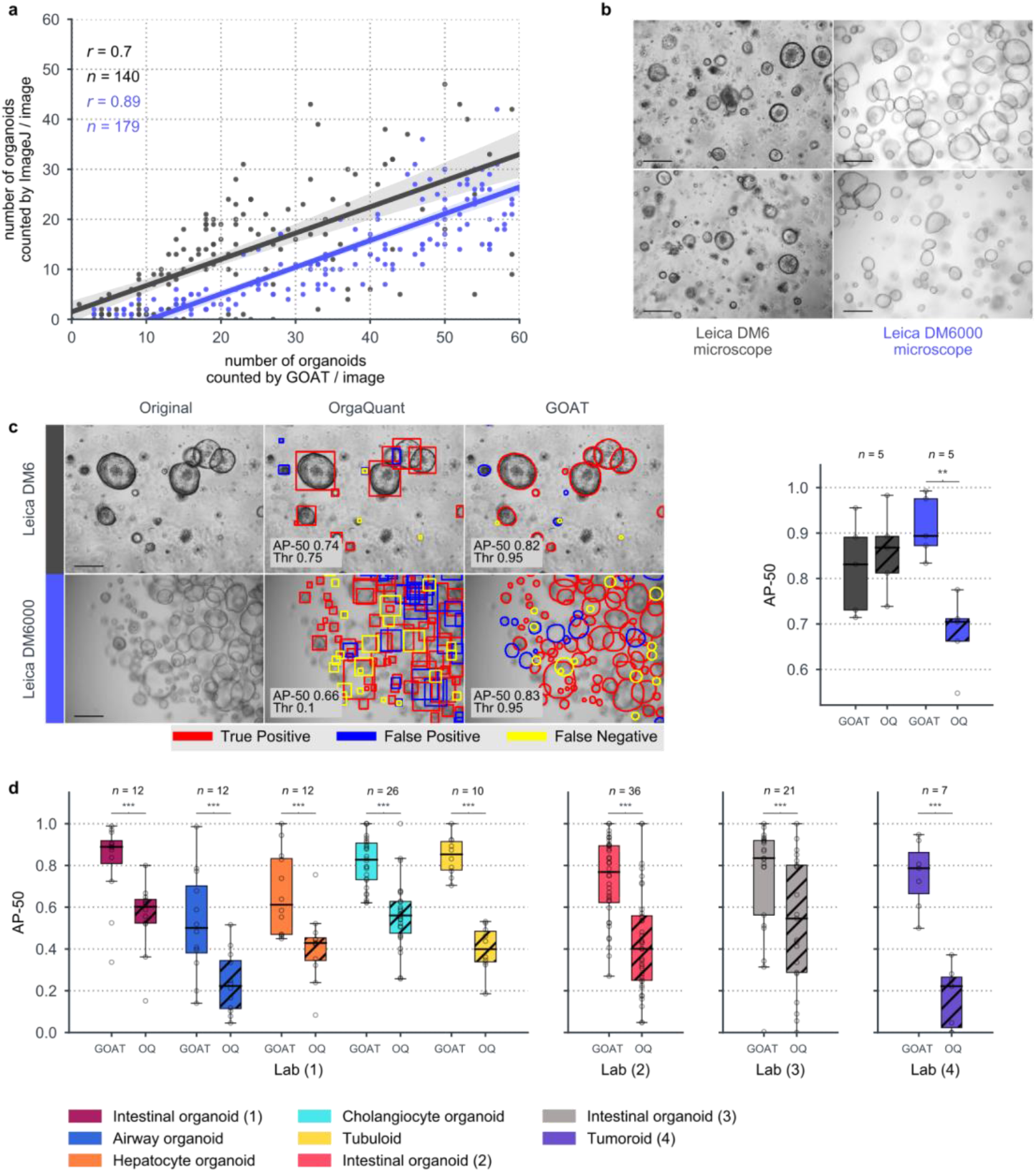
Comparison of organoid annotation with GOAT and other methods. **a**, Correlations between number of organoids in microscopy bright field images predicted by GOAT and manually counted by a researcher using ImageJ, showing a strong correlation between predictions of GOAT and manual annotation (See Methods for details). Images were derived from a previous publication (gut development (GD) dataset). All scale bars represent 250 μm. **b**, Microscopy images of intestinal organoids generated in Lab (2) with two different microscopes, a Leica DM6 and a Leica DM6000 microscope, showing variation in brightness. **c**, Images showing organoid annotation with the previously published OrgaQuant (OQ) method (based on bounding object detection) and GOAT (based on instance segmentation), showing matches (True Positive), missed predictions (False Negative) and additional predictions (False Positive) calculated at a IoU threshold of 0.5 using the GD dataset (see Methods). Comparisons of GOAT and OrgaQuant AP-50 values generated from images of the GD data set imaged with Leica DM6 (gray) and a Leica DM6000 (blue) are shown. For visualization purposes the confidence thresholds (Thr) for the representative images were chosen to have the highest F1-50 score and is indicated in the lower left corner together with the AP-50, which was calculated without setting a confidence threshold. **d)** Comparisons of AP-50 scores generated by GOAT and OrgaQuant on the testing dataset, which includes intestinal, hepatocyte, airway, and cholangiocyte organoids, tubuloids and tumoroids. Medians with interquartile ranges are shown. Pairwise *t*-test \*\**P* < 0.01, \*\*\**P* < 0.001.

Recently, OrgaQuant, a neural network driven approach to quantify organoids in images was developed^10^. OrgaQuant utilizes bounding boxes to annotate instances of organoids in brightfield images, which allows for the quantification of the number of organoids but not advanced organoid features such as size, density and eccentricity. The performances of OrgaQuant and GOAT were compared using a subset of the GD dataset (*n* = 10 images), which was labeled with ellipses by external researchers **(Fig. 3c)**, and our testing dataset (*n* = 136 images) **(Fig. 3d, Supplementary Fig. 2)**. In contrast to the more precise instance segmentation approach used by GOAT, OrgaQuant utilizes a bounding box approach. As OrgaQuant had not determined a confidence threshold for the tool, both models were evaluated on a confidence threshold free metric, the commonly used average precision at an IoU threshold of 0.5 (AP-50 score) (for details see Methods). The AP-50 ranges from one (the predicted masks (PEs) with the highest confidence perfectly match the ground truth masks (GTs)) to zero (no PE matches the GT at any confidence level). Low AP-50 scores are observed upon high confidence FPs. GOAT performed significantly better on microscope 2 in the GD dataset in comparison to OrgaQuant with AP-50 scores of 0.91 and 0.68 respectively (t(9) = 6.81, *P* < 0.01). The performance of GOATs and OrgaQuants analyses of images taken with microscope 1 did not differ (AP-50 GOAT=0.82 and AP-50 OrgaQuant=0.86; t(9) = −0.85, *P* = 0.59). Next, the performance of GOAT and OrgaQuant to annotate organoids derived from other tissues were compared. GOAT outperformed OrgaQuant on all tissues, except intestinal lab (3), included in our testing dataset (AP-50 values Lab (1): intestinal organoids GOAT: 0.82 vs OrgaQuant: 0.56 (t(23) = 8.49, *P* < 0.001), airway organoids: 0.52 vs 0.24 (t(23) = 5.39, *P* < 0.001), hepatic organoids: 0.67 vs 0.41 (t(23) = 5.52, *P* < 0.001), cholangiocyte organoids: 0.82 vs 0.57 (t(51) = 9.31, *P* < 0.001), tubuloids 0.85 vs 0.40 (t(19) = 17.03, *P* < 0.001); Lab (2): intestinal organoids: 0.73 vs 0.43 (t(71) = 7.16, *P* < 0.001); Lab (3): intestinal organoids: 0.74 vs 0.53 (t(41) = 4.03, *P* < 0.001), tumoroids: 0.76 vs 0.17 (t(13) = 7.81, *P* < 0.001)) (**Fig. 3d)**. Very recently, another deep learning based tool for organoid quantification was published on bioRxiv^24^. This approach is based on semantic segmentation **(Fig. 1c)**, which does not discriminate between overlapping organoids in the same image nor quantifies advanced features such as eccentricity and density. Taken together, GOAT is the only tool available to provide pixel-level precise annotation of overlapping organoids by design and performs significantly better compared to other available tools for organoid annotation especially on more diverse organoid cultures such as tumoroids and cholangiocyte organoids while allowing for comparative studies between different laboratories and imaging techniques.

## Discussion

Organoid systems are increasingly used in fundamental and clinical research; however, fast and robust annotation methods are lacking. Here, we present the development of GOAT, a neural network-driven tool for organoid annotation. GOAT showed high performance and generalization for multiple organoid systems generated from different tissues, imaging tools and laboratories. The novelty of GOAT, compared to previous approaches, lies in its instance segmentation approach enabling precise pixel-level annotation and phenotyping of organoids. These quantifications were not possible with previous methods using rudimentary bounding boxes lacking important pixel-level precision and semantic segmentation, as overlapping organoids could not be identified. Annotating microscopy images with GOAT resulted in a time gain from weeks of manual annotation to minutes of automatic analysis, making high-throughput and real-time analyses of a diverse range of organoid systems a reality.

Even though we evaluated GOAT on a wide range of organoids and experimental settings, the model may find its limitations in rare organoid shapes, such as tumoroids, which may result in poorer performance. Furthermore, backgrounds of organoid images are frequently covered by other tissue or immune cells, such as airway organoid cultures cultivated and seeded from bronchoalveolar lavage samples, possibly reducing quantification performance. Further training of the neural network on additional datasets, including heterogeneous cell populations, will remedy these limitations and improve the versatility of GOAT.

Taken together, we present a fast, high-throughput analysis tool to phenotype multiple organoid systems with pixel-level precision, using deep neural networks to achieve high accuracy of organoid annotation. GOAT can now be used to perform robust and comparative studies employing organoids by quantifying growth kinetics and deeper phenotyping of organoids to accelerate organoid research and pave the way for high-throughput studies and real-time assessment of patient specific organoids.

## Code Availability

The code base for GOAT is available at: https://github.com/msknorr/goat-public

## Data availability

The imaging data, as well as the instance segmentation masks that support the findings will be made available at publication at www.zenodo.com. The datasets acquired from external laboratories should be directly requested from these.

## Methods

### Annotation procedure

In the first phase of annotation, intestinal organoids were labeled using bounding boxes in brightfield microscopy images (*n* = 246), which then were used to train an object detection model. This object detection model was used to predict pseudo labels on the remaining unlabeled data. These pseudo labels were then revised by the researchers, who edited, removed and added bounding boxes where appropriate. This procedure was used to label images of intestinal organoids from 3 laboratories (Lab (1): *n* = 83 images, Lab (2): *n* = 36, Lab (3): *n* = 21), organoids from other tissues from Lab (1) including hepatic (*n* = 94), airway (*n* = 206), and cholangiocyte organoids (*n* = 26) and tubuloids (*n* = 10) as well as CRC tumoroids (*n* = 7) from Lab (4). Similarly, in the second phase, organoid bounding boxes (*n* = 300) were cropped and a segmentation mask was manually drawn on them. The crops were used to train a semantic segmentation model. This semantic segmentation model predicted pseudo labels, which then were manually edited where needed.

### Models

In sum, three different models were employed in this project. (1) GOAT: the final Mask R-CNN instance segmentation model, (2) an Efficient-Det object detector for accelerating the labeling process by generating pseudo labels, (3) a U-Net model for facilitating the generation of instance segmentation masks from bounding box-annotated images.

### GOAT: Instance Segmentation

GOAT consists of a Mask R-CNN architecture with a ResNet-34-FPN backbone. For training and inference, images were resized to 512 × 512 shape. The learning rate was initialized with 0.0001, and reduced by a factor of 0.5 after 5 epochs without performance improvement on the validation set. The Adam optimizer was used. The ResNet34-backbone utilized COCO-dataset^26^ pretrained weights. The batch size was 3 and the region of interest batch size was set to 256. The total loss consists of the equally weighted sum of 5 single losses: (1) The Box Regression Loss, (2) Mask Loss, (3) Classification Loss, (4) Objectness Loss, (5) RPN Box Regression Loss as described in the torchvision github repository. Training augmentations of random hue and saturation shifts, brightness and contrast changes, random rotations and horizontal and vertical flips were applied.

### EfficientDet-D5: Bounding Box Pseudo Labeling

For pseudo labeling of bounding boxes we trained an EfficientDet-D5^27^ with an EfficientNet-B5^28^ backbone. For training and inference, images were resized to 512 × 512 shape. As of the time of this study, full model pre-trained weights for EfficientDet’s have not been released for Pytorch. Therefore, only the backbone was initialized with Image-Net pretrained weights. All other weights were initialized randomly.

### EfficientNet-B0 U-Net: Semantic Segmentation

The segmentation model consists of an U-Net architecture^14^ with an EfficientNet-B0^28^ backbone. Bounding boxes from the original image were cropped and resized to 128 × 128 image size. An initial learning rate of 0.0004 and a batch size of 16 was used. The model was trained with dice loss.

### Performance metrics

#### F1-50

The F1-score is a combined estimate of precision and recall, and is calculated as:

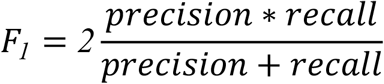

In order to calculate the F1-score in an instance segmentation task first, a confidence threshold that excludes instance segmentation masks of low confidence scores needs to be assigned. The confidence threshold was chosen by computing the F1 score for incremental threshold values (steps of 0.05) on the threshold calculation dataset (*n* = 12). The highest F1-50 score of 0.760 was detected at a confidence threshold of 0.9 and subsequently used for all further analysis **(Fig. 2b)**. As a second threshold, the IoU threshold was determined. The IoU measures the amount of overlap of a predicted instance segmentation mask (PE) and the ground truth instance segmentation mask (GT) in terms of area, as:

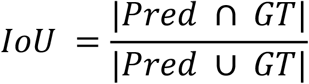

IoU ranges from one (PE and GT pixels match perfectly) to zero (no pixels align). Every PE was paired to every GT and for every pair the IoU was calculated. If a pair’s IoU exceeded the IoU threshold, it was a match (true positive (TP)), starting with the PE with the highest confidence. Each GT can be only assigned to one PE. PEs not matched were determined as false positives (FP) and GTs not matched as false negatives (FN). This allowed determining precision and recall for the F1-calculation, as:

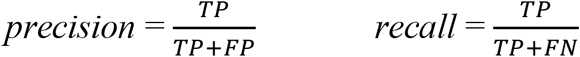

Although IoU thresholds used for machine learning benchmarks vary between publications^22,29,30^, 0.50 as applied here, is most commonly used^23^. To validate this the F1-scores (F1-10, …, F1-90) for different IoU thresholds were calculated **(Fig. 2c)**.

#### AP-50

As OrgaQuant had not determined a confidence threshold for the tool, both models were evaluated on a confidence threshold free metric, the commonly used average precision at an IoU threshold of 0.5 (AP-50 score). Here, masks with low confidences were not removed. TP, FP & FN for each PE were calculated as described above for the F1-50 calculation. Then, the PEs were sorted by confidence and for each PE, precision and recall were calculated from cumulative TP, FN & FP values. From this, a precision recall curve (p(r)) was derived. The Area under the p(r) is the AP-50 score:

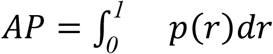

### Quantification metrics

The **size** metric is the median size of all organoids in an image, including segmentation masks on the borders. The **total area** of organoids is the sum of the size of all individual organoids in an image.

The **density (d)** of an organoid is calculated from the mean pixel value of an instance mask, ranging from 0 to 255, which is than normalized as:

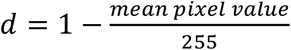

Consequently the metric ranges from 0 to 1. Higher values correspond to a darker, more differentiated organoid and lower values to a brighter, less differentiated organoid^2^.

**Eccentricity (e)** is a measurement for the roundness of a conic section. An ellipse is fitted on the instance mask of each organoid using least square minimization. From this ellipse the semi-major axis (a) and semi-minor axis (b) are derived and eccentricity calculated as:

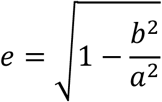

The eccentricity of a parabola is 1. The eccentricity of a circle is 0. The eccentricity of an ellipse is between 0 and 1 with higher values being less circular.

### Statistical Analysis

The comparison of OrgaQuant and GOAT was performed using two tailed paired t-tests. Pearson correlation coefficients were calculated using the SciPy library and default settings.

### Software

The manual annotation of bounding boxes was performed using the VGG Image Annotator^31^. Segmentation annotation was performed using SuperAnnotate^32^. Rechecking of instance segmentation masks was performed in VGG Image Annotator^31^. Model training was performed in Python 3^33^ using PyTorch^34^ and torchvision^35^. All statistical analyses were performed using Python 3^33^ and SciPy^36^. Plots and graphics were generated using Matplotlib^37^.

### Datasets

#### Human Organoids

Human samples were obtained with written informed consent provided by the donors. Tissue samples were obtained by the University Medical Center Hamburg-Eppendorf (UKE), the Asklepios Clinic Hamburg-Barmbek and human immune system Mouse (HIS Mouse) Facility of the Amsterdam Medical Center (AUMC) and the Utrecht Medical Center as reported previously^18,38,39^. Collection of tissue and blood samples was approved by the local ethics committees.

To develop a generalized organoid annotation tool we trained and validated the algorithm on adult stem cell-derived organoids from multiple different tissues (intestine, liver colorectal cancer (CRC)) or body fluids (bronchoalveolar lavage and urine). Generation and culturing of the organoid system is described below.

##### Intestinal organoids

Human intestinal organoids were generated from fetal or adult intestinal tissue samples as previously described^2,6,18,38,39^. Briefly, the epithelial layer was detached from intestines using ethylenediaminetetraacetic acid (EDTA; Sigma-Aldrich, 03690) and 1,4-dithiothreitol (DTT; Carl Roth GmbH+Co.KG, 6908.1; Sigma-Aldrich, D8255) for 1 hr at 4°C. Isolated epithelial cells were washed with AD+++ (Advanced DMEM/F12 (Thermo Fisher, 12634-028) supplemented with 1% GlutaMAX (Thermo Fisher, 35050061), 10 mM HEPES (Thermo Fisher, 15630056; Sigma-Aldrich, H3375), and 1% penicillin/streptomycin (Thermo Fisher, 15140-122; Sigma-Aldrich, P4333)) and resuspended in ice-cold growth factor-reduced Matrigel (Corning, 356231) diluted with AD+++. Three Matrigel droplets of 10 μl containing ISCs were seeded per well in pre-warmed 24-well plates (Greiner, 662160), solidified for 10 minutes at 37°C and covered with 0.5 ml expansion medium (EM; **Supplementary Table 3**) supplemented with 10 μM Rho k inhibitor Y-27632 (Y; StemCell, 72308). EM + Y was refreshed every 2-3 days for 10-14 days until first passage. Intestinal organoid cultures were passaged weekly by mechanical disruption. Organoid fragments were resuspended in ice-cold growth factor-reduced Matrigel diluted with AD+++. Matrigel droplets containing organoid fragments were re-seeded in pre-warmed 24-well plates and covered with 0.5 ml EM. EM was refreshed every 2-3 days. Images of intestinal organoids used for training and validation of GOAT were derived from different experimental setups in multiple laboratories:

1. Fetal intestinal organoids were generated after seeding single cells, cultured in EM + Y or EM + Y supplemented with increasing concentrations of TNF and imaged at day 12 with a Leica DM6 or DM6000 inverted microscope at the Amsterdam University Medical Center in Amsterdam (data published in Schreurs et al., 2019, Immunity)^18^.
2. Adult intestinal organoids were grown from single cells and cultured in complete or reduced EM either supplemented with Wnt3a conditioned medium (WCM, home-made) or Wnt surrogate (U-Proteintech Express B.V., N001-0.1mg). In reduced EM, SB202190 (Sigma-Aldrich, S7067-25MG), A83-01 (Tocris, 2939/10) and nicotinamide (Sigma-Aldrich, N0636-100G) were removed from the complete medium either individually or in combinations. Y was added to all EM variations for the first 2-3 days after seeding the single cells. Images were acquired at day 7, 9 and 12 with an Evos M5000 imaging system (Thermo Fisher) at the Leibniz Institute of Virology in Hamburg (unpublished data).
3. Adult intestinal organoids were grown from single cells in the absence or presence of bead isolated CD3-positive T cells from peripheral blood in complete or reduced EM supplemented with WCM. T cells were either seeded inside the Matrigel or added to the organoid medium. Y was added to all EM variations for the first 2-3 days after seeding the single cells. Images were acquired at day 7, 10 and 12 with an Evos M5000 imaging system at the Leibniz Institute of Virology in Hamburg (unpublished data).
4. Adult intestinal organoids were grown from single cells, cultured in EM (+ Y for the first 2-3 days after seeding) in the absence and presence of different concentrations of EGF, HGF, gefitinib and/or savolitinib. Images were acquired at day 7, 9 and 12 with an Evos M5000 imaging system or an Olympus microscope with a CoolSnap camera at the Amsterdam University Medical Center in Amsterdam (data previously published)^38,39^.

##### Airway organoids

Human airway organoids (AO) were generated from bronchoalveolar lavage fluid (BALs) according to an adapted protocol from Sachs et al^5^. In brief, BALs were incubated with 100 mg/ml primocin (Invivogen, ant-pm-1), 40 mg/ml gentamycin (Thermo Fisher, R01510) and 1 mg/ml amphotericin B (Thermo Fisher, R01510) for 30 min on ice to reduce the risk of bacterial or fungal contaminations. Subsequently, BALs were digested in airway organoid expansion medium (AOEM, **Supplementary Table 4**) supplemented with 0.5 mg/ml Collagenase D (Sigma, 11088882001), 0.5% Sputolysin (Boehringer Ingelheim Vetmedica, SPUT0001), 40 mg/ml gentamycin and 1 mg/ml amphotericin B for 30 min at 37°C. After digestion the cell suspension was filtered through a 100 μm cell strainer (Greiner, 542000). Isolated single cells were washed with AD+++ and resuspended in an appropriate volume of ice-cold BME type II (R&D Systems, 3533-010-02). Two 20 μl droplets of BME type II containing airway epithelial cells were seeded per well in pre-warmed 24-well plates (Greiner, 662160). After solidification of BME type II droplets, 0.5 ml AOEM supplemented with 10 mg/ml gentamycin and 250 ng/ml amphotericin B was added to each well. AOEM supplemented with gentamycin and amphotericin B was refreshed every 3-4 days until first passage. Airway organoid cultures were passaged every 2-3 weeks through mild treatment with TrypLE Express (Thermo Fisher, 12605028). Airway organoid fragments were resuspended in ice-cold BME type II and BME II and 20 μl droplets were seeded in pre-warmed 24-well plates. After solidification of the droplets, 0.5 ml AOEM was added to each well. AOEM was refreshed every 3-4 days. Microscopy images documenting airway organoid growth were acquired with an Evos M5000 imaging system at the Leibniz Institute of Virology in Hamburg.

##### Tubuloids

Tubuloids were generated according to an adapted protocol by Schutgens et al^20^. Approximately 250-300 ml urine was processed directly after voiding. 1% penicillin/streptomycin (Sigma-Aldrich, P4333), 10 μM Rho-kinase inhibitor Y-27632 (StemCell, 72308) and 0.1 mg/ml primocin (Invivogen, ant-pm-1) were added. After centrifugation for 10 min at 500 g and 4°C, the pellet was washed with AD+++, supplemented with 10 μM Y and 0.1 mg/ml primocin. After a second centrifugation step, the pellet was resuspended in ice-cold Matrigel (Corning, 356231). Three Matrigel droplets of 10 μl were seeded per well in pre-warmed 24-well plates (Greiner, 662160). After gelation, 0.5 ml tubuloid culture medium (**Supplementary Table 5**) was added per well. Tubuloids were passaged every 1-2 weeks by mechanical shearing or TrypLE Express (Thermo Fisher, 12605028) digest. Tubuloid fragments were embedded in ice-cold Matrigel diluted with AD+++ and three droplets of 10 μl were seeded per well of pre-warmed 24-well plates. After complete gelation, 0.5 ml tubuloid culture medium was added to each well. Medium was refreshed every 2-3 days. Tubuloid images were acquired with an Evos M5000 imaging system at the Leibniz Institute of Virology in Hamburg.

##### Hepatocyte organoids

Human hepatocyte organoids were generated from adult human liver tissue according to Hu et al^4^. In brief, human liver samples were digested using 2.5 mg/ml Collagenase D (Sigma-Aldrich, 11088882001) and 0.1 mg/ml DNaseI (StemCell, 07470). Subsequently, cells were filtered through a 70 μm cell strainer and washed with wash medium (DMEM high glucose Glutamax (Thermo Fisher, 12077549), 1% FBS (Biochrom AG, S0615), 1% penicillin/streptomycin (Sigma-Aldrich, P4333)). Cells were resuspended in ice-cold BME-type II (R&D Systems, 3533-010-02) and seeded in pre-warmed 24-well plates (Greiner, 662160). After solidification, 0.5 ml hepatocyte organoid medium (**Supplementary Table 6**) was added per well. Medium was refreshed every 2-3 days. Hepatocyte organoids were passaged every week by mechanical disruption. Organoid fragments were embedded in cold hepatocyte organoid medium and BME type II (with a ratio of 1:3) and one 50 μl droplet was seeded per well of pre-warmed 24-well plates. After complete gelation, 0.5 ml hepatocyte organoid medium was added to each well. Medium was refreshed every 2-3 days. Hepatocyte organoid images were acquired with an Evos M5000 imaging system at the Leibniz Institute of Virology in Hamburg.

##### Cholangiocyte organoids

Human cholangiocyte organoids were generated from adult human liver tissues as previously describe^19^ Briefly, minced human liver samples were incubated in a pre-warmed digestion mix containing 2.5 mg/ml Collagenase D (Sigma-Aldrich, 11088882001) and 0.1 mg/ml DNaseI (StemCell, 07470) for 30-90 min at 37°C. Digested material was filtered through a 70 μm cell strainer and washed with ice-cold wash medium (DMEM high glucose Glutamax (Thermo Fisher, 12077549), 1% FBS (Biochrom AG, S0615), 1% penicillin/streptomycin (Sigma-Aldrich, P4333)). The desired number of cells was resuspended in ice-cold BME type II (R&D Systems, 3533-010-02) and one 50 μl droplet was seeded per well in a pre-warmed 24-well plate (Greiner, 662160). After gelation, 0.5 ml cholangiocyte organoid isolation medium (**Supplementary Table 7**) was added per well. The isolation medium was exchanged 3-4 days after seeding by cholangiocyte organoid expansion medium (**Supplementary table 8**). Cholangiocyte organoid expansion medium was refreshed every 3-4 days. Cholangiocyte organoid cultures were passaged weekly by mechanical disruption. Cholangiocyte organoid fragments were resuspended in cold BME type II and 50 μl droplets were seeded per well of pre-warmed 24-well plates. After polymerization of the basement matrix, each well was covered with 0.5 ml cholangiocyte organoid expansion medium. Medium was refreshed every 3-4 days. Cholangiocyte organoid images were acquired with a Nikon A1 microscope at the Leibniz Institute of Virology in Hamburg.

##### CRC tumoroids

Human colorectal cancer (CRC) tumoroids were generated from tissue samples of treatment naive CRC patients^2,6,40^. Minced tumor tissue was incubated on a rocking plate for 30 min at 37°C with Gentle Cell Dissociation Reagent (STEMCELL, 07174). Digested tissue was washed with ice-cold PBS with 0.05% BSA and filtered (>35 μm and <70 μm Easy Strainer). Crypts were resuspended in DMEM/F-12 (Gibco, 11554546) with 15 mM HEPES (Gibco, 11560496) and Matrigel (Corning, 356231) in a 1:1 ratio. One 50 μl droplet was seeded per well in a pre-warmed 24-well plate (Corning, CLS3527-100EA). After gelation, 0.75 ml StemPro medium (Thermo Fisher, A1000701) was added to each well. CRC tumoroids were passaged every two weeks after mechanical disruption. Tumoroid fragments were resuspended in a 1:1 mix of ice-cold Matrigel and StemPro medium and seeded in pre-warmed 24-well plates. Each well was covered with 0.75 ml StemPro medium after solidification of the Matrigel droplets. Medium was refreshed every five days. CRC tumoroid images were acquired using the cell imager Paula (Leica) at the University Medical-Center Hamburg-Eppendorf.

## Acknowledgements

This work was supported by the the Free and Hanseatic City of Hamburg (Landesforschungsförderung Hamburg: LFF-FV73, LFF-FV75, LFF-FV78), Deutsche Forschungsgemeinschaft (BU 3630/2-1), Deutsche Zentrum für Infektionsforschung, EFRE 2014-202 REACT-EU, Dutch Digestive Fund (MLDS CDG 15-02), the Deutsche Forschungsgemeinschaft (DFG; SFB841) and the Daisy Huët Röell Foundation. The Leibniz Institute of Virology is supported by the Free and Hanseatic City of Hamburg and the Federal Ministry of Health.

## Author information

### Contributions

J.P.B. and M.S.K. designed and developed the software for all the proposed computational analyses. M.E.B., J.P.B., M.S.K., M.A. and M.J.B designed the experiments. M.E.B., L.W., J.W, A.J.P., J.M.J, A.J.H., J.J., O.H., S.B, J.N., L.M.M., A.N., L.M., S.L., T.K., S.P., N.G., R.R.C.C.S., S.J. and L.K. contributed to experiments and organoid image annotation. F.W.R.V, K.O., C.S. D.P., K.R., I.K. and S.K. collected tissue samples. J.P.B. and M.S.K. analyzed the data. J.P.B., M.S.K., M.T., M.E.B. and M.J.B. wrote the paper with input and comments from all authors. M.J.B. and M.A. conceived and supervised the project.

### Ethics declarations

Human samples were obtained with written informed consent provided by the donors. Tissue samples were obtained by the University Medical Center Hamburg-Eppendorf (UKE), the Asklepios Clinic Hamburg-Barmbek and human immune system Mouse (HIS Mouse) Facility of the Amsterdam Medical Center (AUMC) and the Utrecht Medical Center as reported previously^18,38,39^. Collection of tissue samples was approved by the local ethics committees.

## Competing interests

J.P.B., M.S.K., M.J.B. and M.A. are inventors of one provisional patent that describe the methods in this paper.

