## Supplemental Information for "GOAT: Deep learning-enhanced Generalized Organoid Annotation Tool"

### Supplementary Information

#### Supplementary Fig. 1: Alternative performance metrics for GOAT.

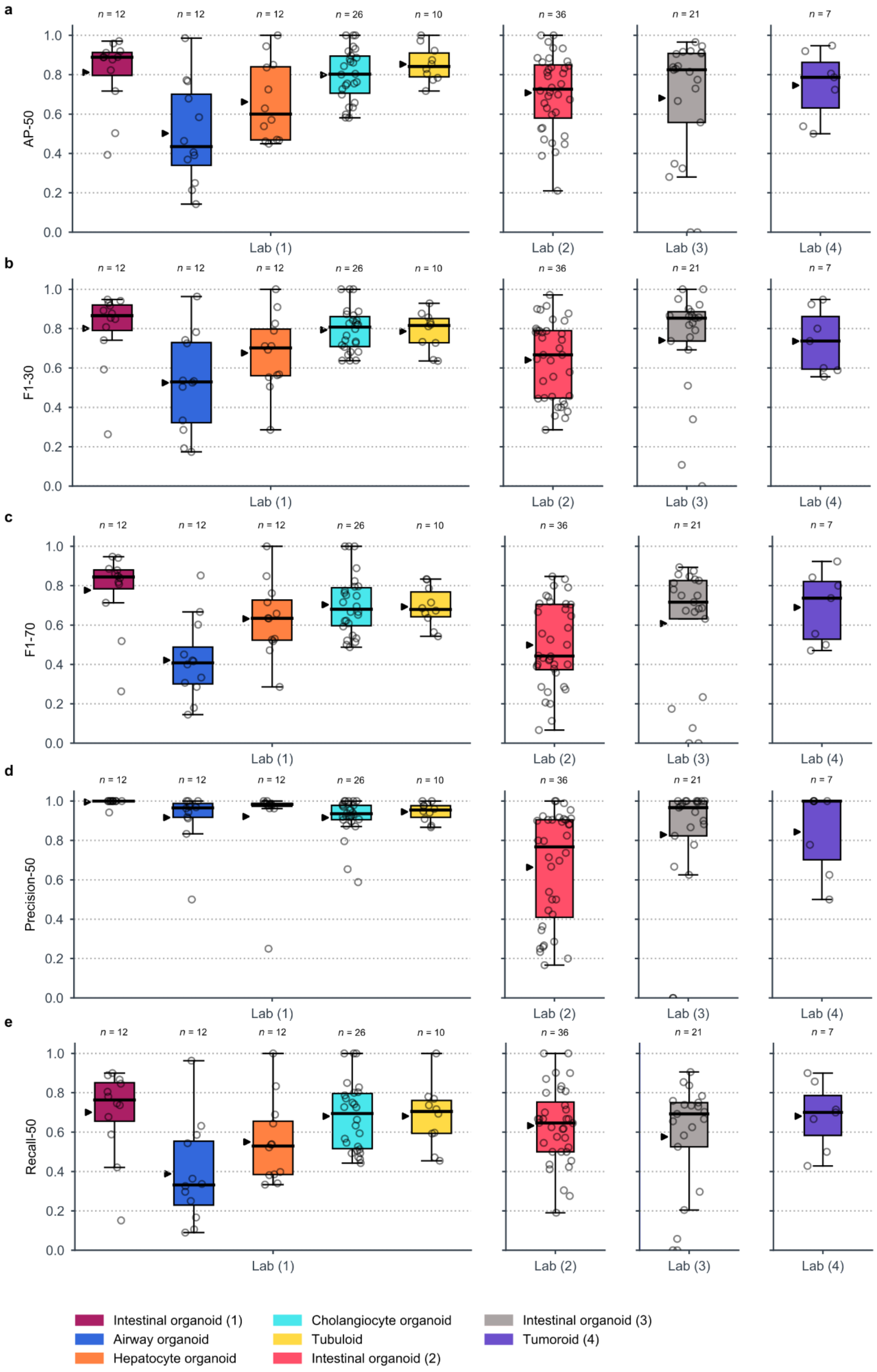

**a**-**e,** Alternative performance metrics on the testing dataset for intestinal, cholangiocyte, airway, hepatocyte organoids, tubuloids and tumoroids including AP-50 (**a**) common metric to validate instance segmentation tasks, (**b**) F1-30 and (**c**) F1-70 are alternatives to F1-50 using different IoU thresholds, (**d**) precision at a confidence threshold of 0.9 for obtaining the F1-50 score and (**e**) recall at a confidence threshold of 0.9 utilized for the F1-50 score.

**Supplementary Fig. 2: Performance examples of GOAT and OrgaQuant on the test dataset.**

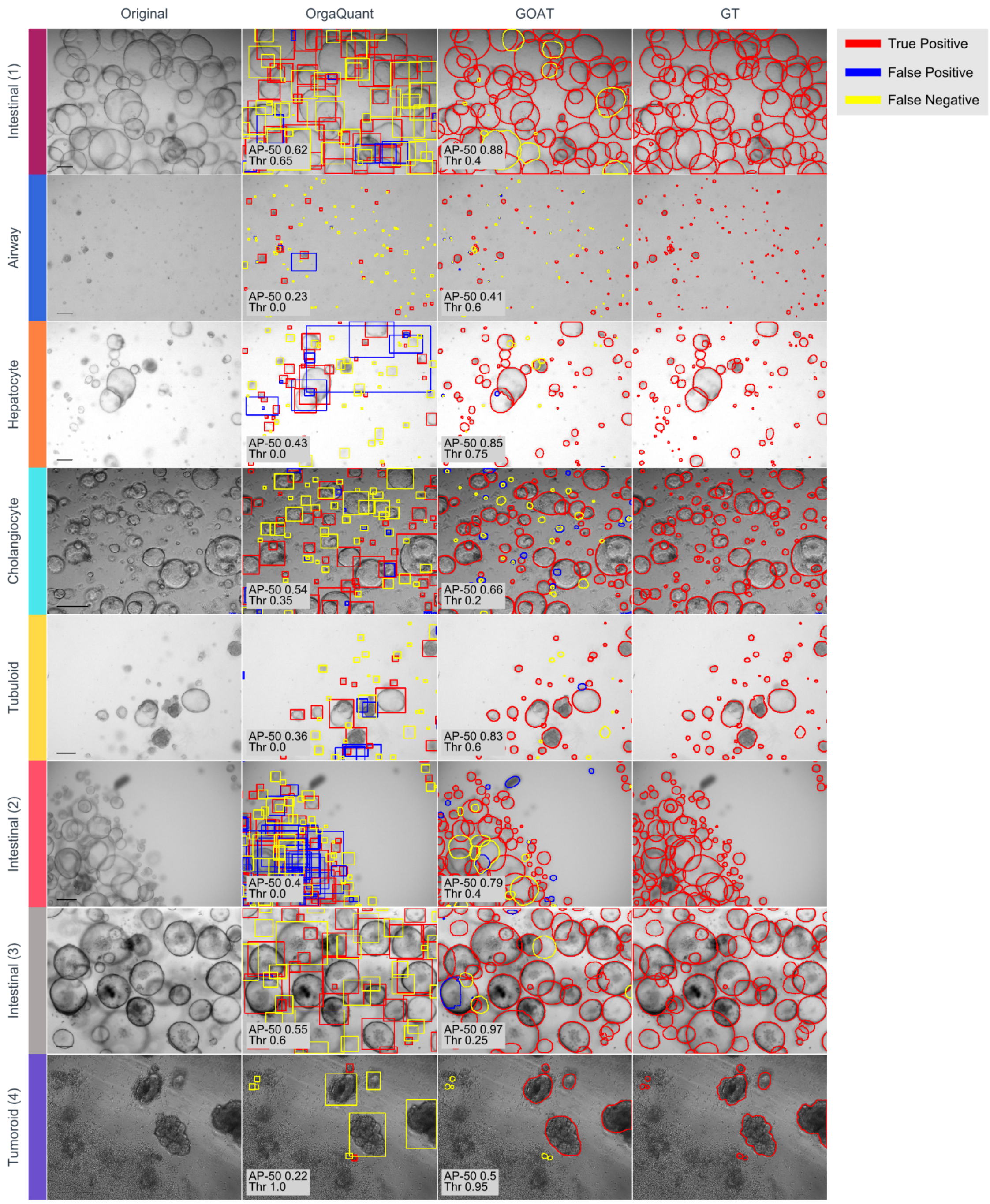

Comparison of the performance of GOAT using instance masks and OrgaQuant using bounding boxes at IoU threshold of 0.5 analyzing the test dataset. For visualization purposes the confidence thresholds (THR) for the representative images were chosen to have the highest F1-50score. The AP-50 was derived from correct predictions (true positive, TP, red), missed predictions (false negative, yellow) and additional predictions (false positive, blue). Thr for the visualization and AP-50 scores, calculated without setting a threshold, are indicated in the bottom left corner.

**Supplementary table 1**: Number of images per dataset and organoid system

| | HIO lab (1) | AO | Hep Org | CholOrg | Tubuloid | HIO lab (2) | HIO lab (3) | Tumor-oid lab (4) | Total |
|---|---|---|---|---|---|---|---|---|---|
| Training | 281 | 161 | 46 | 0 | 0 | 0 | 0 | 0 | 488 |
| Validation | 32 | 29 | 32 | 0 | 0 | 0 | 0 | 0 | 93 |
| Threshold Calculation | 4 | 4 | 4 | 0 | 0 | 0 | 0 | 0 | 12 |
| Testing | 12 | 12 | 12 | 26 | 10 | 36 | 21 | 7 | 136 |
| Total | 329 | 206 | 94 | 26 | 10 | 36 | 21 | 7 | 729 |

HIO: human intestinal organoids; AO: Airway organoids;HepOrg: hepatocyte organoids; CholOrg: cholangiocyte organoids.

**Supplementary table 2**: Diverse performance metrics for GOAT on testing data

| | HIO lab (1) | AO | Hep Org | CholOrg | Tubuloid | HIO lab (2) | HIO lab (3) | Tumor-oid lab (4) | mean |
|---|---|---|---|---|---|---|---|---|---|
| F1 | 0.8 | 0.51 | 0.67 | 0.77 | 0.78 | 0.60 | 0.66 | 0.74 | 0.69 |
| Precision | 1.00 | 0.92 | 0.92 | 0.92 | 0.95 | 0.66 | 0.83 | 0.84 | 0.88 |
| Recall | 0.70 | 0.39 | 0.55 | 0.68 | 0.68 | 0.63 | 0.58 | 0.68 | 0.61 |
| AP-50 | 0.81 | 0.50 | 0.66 | 0.80 | 0.85 | 0.71 | 0.68 | 0.75 | 0.72 |
| AP-75 | 0.73 | 0.29 | 0.53 | 0.62 | 0.61 | 0.48 | 0.56 | 0.66 | 0.56 |
| mAP-50:95 | 0.65 | 0.29 | 0.48 | 0.55 | 0.57 | 0.45 | 0.51 | 0.54 | 0.50 |

HIO: human intestinal organoids; AO: Airway organoids; HepOrg: hepatocyte organoids; CholOrg: cholangiocyte organoids.

**Supplementary table 3**: Intestinal organoid expansion medium (EM).

| | Final concentration | company | catalogue number |
|---|---|---|---|
| AD+++ | 20% (v/v) | | |
| B27 supplement | 1x | Thermo Fisher | 17504001 |
| N2 supplement | 1x | Thermo Fisher | 17502001 |
| Murine EGF | 50 ng/ml | Peprotech | 315-09-500 |
| N-acetyl-L-cysteine | 1.25 mM | Sigma-Aldrich | A9165-25G |
| [Leu15]-Gastrin | 10 nM | Sigma-Aldrich | G9145-.1MG |
| Nicotinamide | 10 mM | Sigma-Aldrich | N0636-100G |
| SB202190 | 10 µM | Sigma-Aldrich | S7067-25MG |
| A83-01 | 500 nM | Tocris | 2939/10 |
| Noggin-CM<br>or recombinant human Noggin | 10% (v/v)<br>100 ng/mL | home-made<br>Peprotech | <br>120-10C |
| R-spondin-1-CM<br>or recombinant human R-spondin1 | 20% (v/v)<br>500 ng/mL | home-made<br>R&D Systems | <br>4645-RS |
| WNT3a-CM<br>or Wnt3A surrogate | 50% (v/v)<br>0.5 nM | home-made<br>U-Protein Express | <br>N001-0.1mg |

**Supplementary table 4**: Airway organoid expansion medium (AOEM).

| | Final concentration | company | catalogue number |
|---|---|---|---|
| AD+++ | 70% (v/v) | | |
| B27 supplement | 1x | Thermo Fisher | 17504001 |
| N-acetyl-L-cysteine | 1.25 mM | Sigma-Aldrich | A9165-25G |
| Nicotinamide | 5 mM | Sigma-Aldrich | N0636-100G |
| A83-01 | 500 nM | Tocris | 2939/10 |
| FGF-10 | 100 ng/ml | Peprotech | AF-100-26 |
| FGF-7 | 25 ng/ml | Peprotech | 100-19 |
| SB202190 | 0,5 µM | Sigma-Aldrich | S7067-25MG |
| Y-27632 | 5 µM | StemCell | 72308 |
| Primocin | 50 mg/ml | Invivogen | ant-pm-1 |
| Noggin-CM | 10% (v/v) | home-made | |
| R-spondin-1-CM | 20% (v/v) | home-made | |

**Supplementary table 5**: Tubuloid culture medium.

| | Final concentration | company | catalogue number |
|---|---|---|---|
| AD+++ | 50% (v/v) | | |
| B27 supplement | 1.5x | Thermo Fisher | 17504001 |
| N-acetyl-L-cysteine | 1 mM | Sigma-Aldrich | A9165-25G |
| A83-01 | 5 μM | Tocris | 2939/10 |
| FGF-10 | 100 ng/ml | Peprotech | AF-100-26 |
| Human EGF | 50 ng/ml | Thermo Fisher | PHG0315 |
| IL-4 | 2 ng/ml | R&D Systems | 204-IL-020 |
| Y-27632 | 10 μM | StemCell | 72308 |
| Primocin | 0.1 mg/ml | Invivogen | ant-pm-1 |
| R-spondin-1-CM | 10% (v/v) | home-made | |
| WNT3A-CM | 40% (v/v) | home-made | |

**Supplementary table 6**: Hepatocyte organoid medium.

| | Final concentration | company | catalogue number |
|---|---|---|---|
| AD+++ | 81% (v/v) | | |
| B27 (w/o vitamin A) | 2% (v/) | Thermo Fisher | 17504001 |
| N-acetyl-L-cysteine | 1.25 mM | Sigma-Aldrich | A9165-25G |
| Nicotinamide | 10 mM | Sigma-Aldrich | N0636-100G |
| A83-01 | 2 µM | Tocris | 2939/10 |
| FGF-10 | 100 ng/ml | Peprotech | AF-100-26 |
| FGF-7 | 100 ng/ml | Peprotech | 100-19 |
| Human EGF | 50 ng/ml | Peprotech | AF-100-15 |
| Y-27632 | 10 µM | StemCell | 72308 |
| [Leu15]-Gastrin | 10 nM | Sigma-Aldrich | G9145-.1MG |
| CHIR99021 | 3 µM | Tocris | 4423 |
| HGF | 50 ng/ml | Peprotech | 100-39H |
| TGF-a | 20 ng/ml | Peprotech | 100-16A |

| | | |
|---|---|---|
| R-spondin-1-CM | 15% (v/v) | home-made |

**Supplementary table 7**: Cholangiocyte organoid isolation medium.

| | Final concentration | company | catalogue number |
|---|---|---|---|
| AD+++ | 35% (v/v) | | |
| B27 (minus vitamin A) | 2% (v/v) | Thermo Fisher | 17504001 |
| N2 | 1% (v/v) | Thermo Fisher | 17502001 |
| N-acetyl-L-cysteine | 1 mM | Sigma-Aldrich | A9165-25G |
| Nicotinamide | 10 mM | Sigma-Aldrich | N0636-100G |
| A83-01 | 5 μM | Tocris | 2939/10 |
| FGF-10 | 100 ng/ml | Peprotech | AF-100-26 |
| Human EGF | 50 ng/ml | Peprotech | AF-100-15 |
| [Leu15]-Gastrin | 10 nM | Sigma-Aldrich | G9145-.1MG |
| HGF | 25 ng/ml | Peprotech | 100-39H |
| Forskolin | 10 μM | Tocris | 1099 |
| R-spondin-1-CM | 10% (v/v) | home-made | |

| | | | |
|---|---|---|---|
| Y-27632 | 10 μM | StemCell | 72308 |
| Wnt3a-CM | 30% (v/v) | Home-made | |
| Noggin-CM | 20% (v/v) | Home-made | |

**Supplementary table 8**: Cholangiocyte organoid expansion medium.

| | Final concentration | company | catalogue number |
|---|---|---|---|
| AD+++ | 85% (v/v) | | |
| B27 (minus vitamin A) | 2% (v/v) | Thermo Fisher | 17504001 |
| N-2 | 1% (v/v) | Thermo Fisher | 17502001 |
| N-acetyl-L-cysteine | 1 mM | Sigma-Aldrich | A9165-25G |
| Nicotinamide | 10 mM | Sigma-Aldrich | N0636-100G |
| A83-01 | 5 μM | Tocris | 2939/10 |
| FGF-10 | 100 ng/ml | Peprotech | AF-100-26 |
| Human EGF | 50 ng/ml | Peprotech | AF-100-15 |
| [Leu15]-Gastrin | 10 nM | Sigma-Aldrich | G9145-.1MG |
| HGF | 25 ng/ml | Peprotech | 100-39H |
| Forskolin | 10 μM | Tocris | 1099 |
| R-spondin-1-CM | 10% (v/v) | home-made | |
